# Social Curiosity in Monkeys

**DOI:** 10.1101/2022.11.15.516657

**Authors:** Jessica A. Joiner, Amrita R. Nair, Nicholas A. Fagan, Steve W. C. Chang

## Abstract

Humans and nonhuman animals derive value from many different sources. Some of these sources, notably primary reinforcers like food, are clearly rewarding and can powerfully shape our behaviors. Other sources of reward, such as information, are more intangible and abstract. Like humans, nonhuman primates also find information inherently rewarding, displaying a preference to reveal information about upcoming primary reinforcers even though the information has no bearing on the outcome itself. For animals living in social groups, the need for information extends beyond primary reinforcers like food. They need to acquire information from others or about others. This is especially true for animals, such as nonhuman primates, which live in large and often hierarchically organized societies, where processing social information for the purpose of learning and socializing can be just as critical to survival. To quantify curiosity for knowing social information in monkeys, we extend the advanced information-seeking paradigm (Bromberg-Martin and Hikosaka, 2009) into the realm of abstract, social information. We first replicated the finding that monkeys prefer advanced information about juice size (Bromberg-Martin and Hikosaka, 2009). We next trained monkeys on a social variant of this task. In the advanced social information-seeking task, monkeys had the option to choose a cue that tells them in advance the valence, or facial expression, on a monkey’s face that they will be viewing before receiving the invariant amount of juice. Even though this cue did not impact which facial expression the monkeys would see, they preferred to know the valence of the facial expression in advance. Our results indicate that information-seeking behavior generalizes to seeking social information. Our findings also suggest that curiosity in nonhuman primates can be translated into increasingly abstract levels of information.

## Introduction

Learning about the world and making adaptive decisions is a critical feature of cognition that allows human and nonhuman animals to manipulate their environment to survive. Humans and nonhuman animals derive value from many different sources. Primary reinforcers like food are clearly rewarding, but animals also experience rewards from other categories of stimuli. Some of these sources are abstract and intangible, such as advanced information and vicarious reward. Monkeys find earlier receipt of information rewarding, displaying a preference to reveal advanced information about upcoming primary reinforcers even though the information has no bearing on the outcome itself (Bromberg-Martin and Hikosaka, 2009). Similarly, though it does not impact their own reward values or outcomes, animals find watching another animal receive rewards to be reinforcing as well, which is an example of vicarious reward (Bandura et al., 1963; Mobbs et al., 2009; Chang et al., 2011).

Gathering and seeking information is critical to the survival of animals, and so, the inherently rewarding nature of information is highly plausible. Although information is intangible, it has real practical value to any organism with the capacity to make use of it. If an animal needs to eat, it needs to be able to acquire information from its environment about where and when food or other vital resources will be available. However, most animals, and particularly humans, do not simply live in a vacuum of foraging for food and surviving individually. The types of information animals need to access and utilize are multitudinous. In animals living in social groups, acquiring information from others or about others could be tremendously beneficial. This is especially true for animals, such as humans and nonhuman primates, which live in large and often hierarchically organized societies, where processing social information for the purpose of learning and socializing can be just as critical to survival as processing information about the nature of food resources (Mitani et al., 2012; Chang et al., 2013).

Curiosity, or information-seeking behavior, is considered a motivating drive that developed for its evolutionary advantages in foraging. The function of curiosity is, at its source, to motivate learning (Loewenstein, 1994; Kidd and Hayden, 2015). Lowenstein’s information gap theory (Loewenstein, 1994) holds that curiosity functions like other drive states, like hunger, in which the organism performs a behavior to reduce the need for an intrinsic drive. Furthermore, it has been proposed that curiosity evolved into a drive state that is always “hungry” for information regardless of immediate reward consequences (James, 2008). If this were the case, it is likely that information-seeking tendencies also apply to intangible sources, such as social information.

It remains unknown to what extent nonhuman primates are curious about others. If curiosity is something that nonhuman primates can direct towards others, then their drive for social information should manifest as a willingness to work for the information. However, given that curiosity may have evolved for foraging purposes, it is possible that curiosity for social information will be weaker than that for primary reward like food. To understand rudimentary aspects of social curiosity in nonhuman primates (rhesus macaques, *Macaca mulatta*), we developed a novel social advanced information-seeking task, based on the advanced juice size information-seeking task originally developed by Bromberg-Martin and Hikosaka, to examine monkeys’ tendencies to seek advanced social information. In doing so, we also examined their tendencies to seek advanced juice size information for reference.

## Methods

To assess monkeys’ interest in social information, we adapted an existing information-seeking paradigm originally developed by Bromberg-Martin and Hikosaka (Bromberg-Martin and Hikosaka, 2009). Monkeys were first trained on the original juice size information-seeking task (**Fig. 1**), and then given the opportunity to learn our novel social information-seeking task (**Fig. 2**). In both tasks, monkey’s gaze positions (horizontal and vertical) were sampled at 1,000 Hz by an infrared eye-monitoring camera system (SR Research Eyelink). Monkeys faced a display screen on which stimuli were presented. Stimuli were controlled by a computer running MATLAB (Mathworks). Monkeys were fitted with a juice tube for delivering rewards. The solenoid valves that deliver the liquid rewards were placed in another room to prevent the monkeys from forming secondary associations between solenoid clicks and different reward types. Masking white noise was always played in the experimental room. When the monkeys successfully completed a trial, a sound clip was played. When the monkeys failed to complete a trial, an error noise was played. All experiments were carried out in a dimly lit room. The monkeys were head-restrained during the experiments. Two monkeys, a male (monkey 1, M1) and a female (monkey 2, M2), participated in this experiment. They were both 8 years old and weighed 12 kg and 9 kg, respectively. Both monkeys were implanted with a headpost (Grey Matter Research) to allow for stabilization of the head for eye tracking.

**Figure 1.**
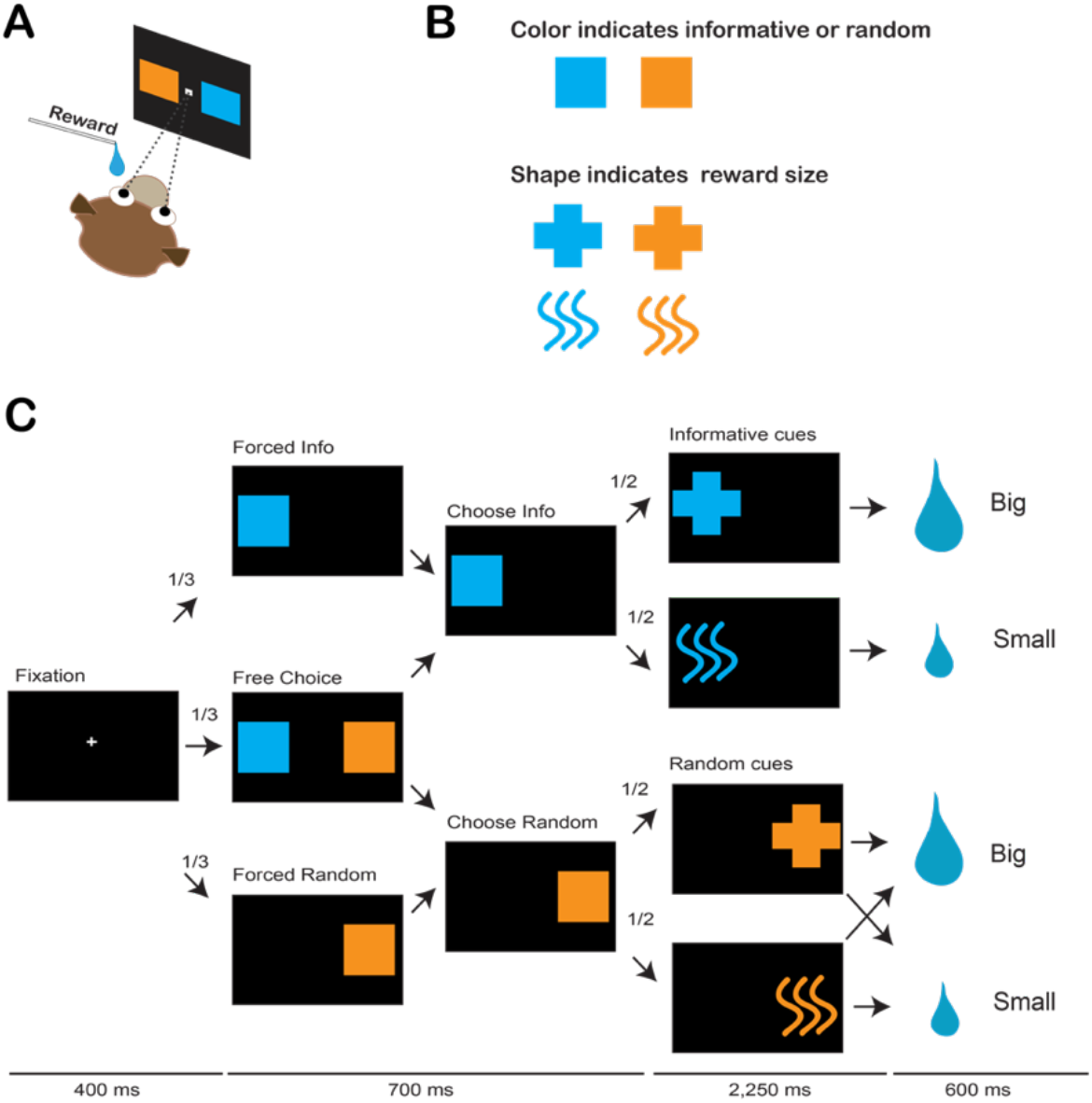
The advanced juice size information-seeking task. **A**. Monkeys sat in a primate chair with a mounted juice tube that delivered juice rewards and faced a monitor to interact with visual stimuli with gaze. **B**. The color of the occluder stimulus (see **C**) indicated if the outcome would be informative or random. The shape of the cue indicated the size of the upcoming juice reward. **C**. The task sequence of the advanced juice size information-seeking task (see Methods for details).

**Figure 2.**
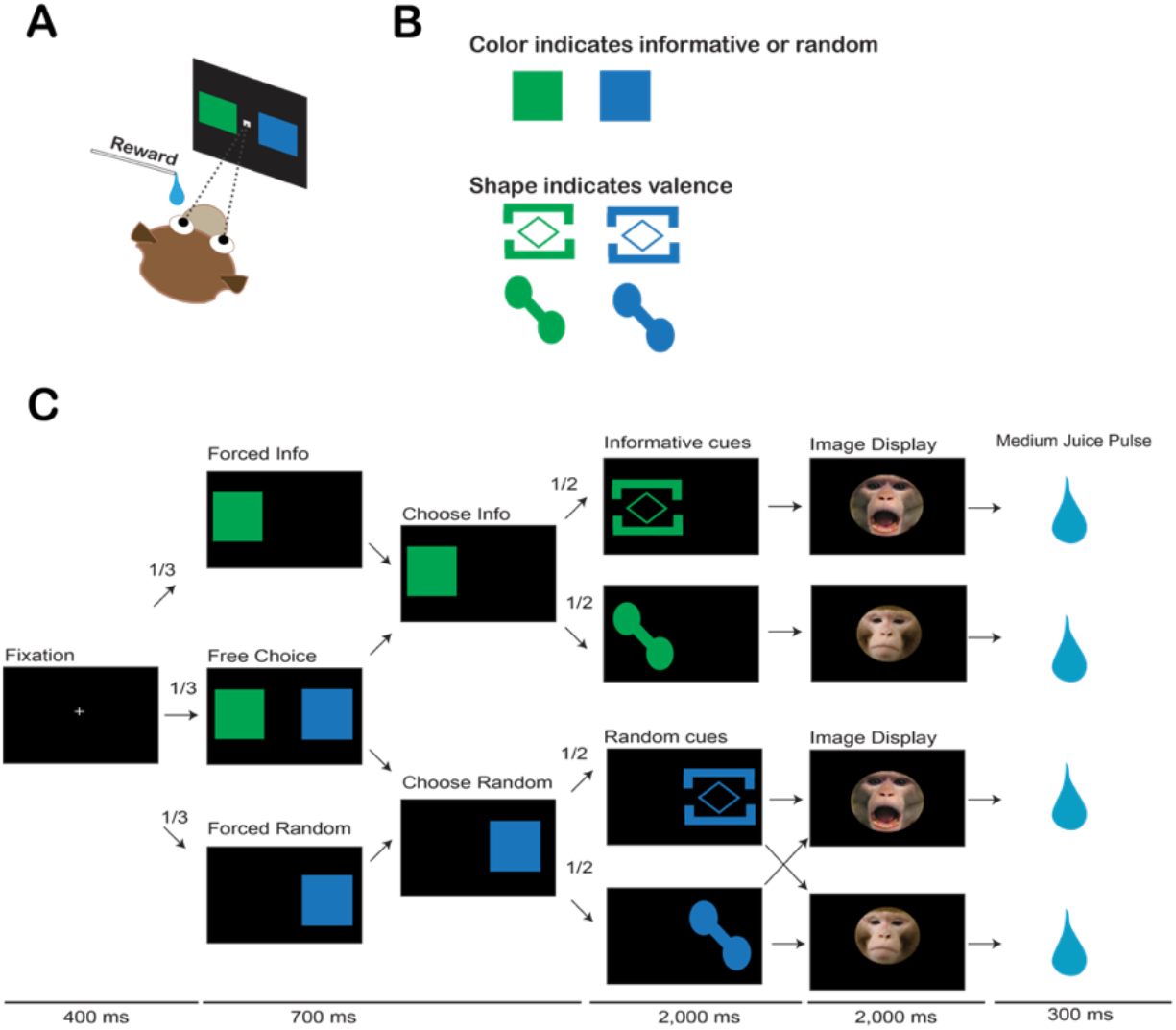
The advanced social information-seeking task. **A**. Monkeys sat in a primate chair with a mounted juice tube that delivered juice rewards and faced a monitor to interact with visual stimuli with gaze. **B**. The color of the occluder stimulus (see **C**) indicated if the outcome would be informative or random. The shape of the cue indicated the size of the upcoming juice reward. **C**. The task sequence of the advanced social information-seeking task (see Methods for details).

### Juice size information-seeking task

In the advanced juice size information-seeking task, monkeys chose to see a cue that reliably tells them how much juice they are about to receive in an environment with variable reward magnitude. However, the choice to receive that information did not alter the amount of juice received. It simply allowed the animal to have advanced information on reward magnitude. This task design was optimal for dissociating reward value and information-seeking, as the monkeys experienced the same reward outcome no matter what. A choice for viewing the information about juice quantity would suggest that the animal would like to know in advance for the sake of having the information.

A critical feature of the task design is that the timing of each task stage was carefully controlled to remove any time-dependent effects on behaviors (**Fig. 1**). This ensures that differing amounts of time did not elapse between each trial depending on the animal’s response. As reward delay is thought to impact the value of the reward (Frederick et al., 2002; Hayden and Platt, 2007), time to reward remained invariant across each trial.

After monkeys initiated a trial by fixating on a central stimulus for 400ms (**Fig. 1**), they were presented with one of three trial types: *forced information, forced random*, and *free choice*. On each of these trials, two occluders appeared, where the color of the occluder indicated if choosing it will lead to an informative cue, or a non-informative, random cue (**Fig. 1**). Monkeys learned the stimulus mapping by trial and error. On forced information and forced random trials, only one occluder appeared, and monkeys were required to fixate on it until the 700ms “state” had elapsed. On free choice trials, monkeys could freely choose between the informative and the random cue. Fixation times were variable, as the choice state was fixed at 700ms, but a minimum fixation of 150ms was required. If monkeys failed to complete the fixation for the entire duration, they were presented with a correction trial. On these correction trials, only the occluder they partially fixated on during the previously aborted trial was presented. Correction trials were not included in any analyses and only existed to help the monkeys learn.

After completing the fixation on the occluder, the monkeys were no longer required to fixate for the remainder of the trial. However, a new cue appeared on the screen in the same location as the fixated occluder. If the monkeys had selected to reveal information before, the shape of the cue accurately predicted the size of the upcoming reward. If they had chosen the random cue, the shapes of the cues were crossed with reward size so that it was correctly predicting the size 50% of the time. This cue remained on the screen for 2,250ms, and then the monkeys entered the reward period. During the reward period, the monkeys were presented with a black screen for 600ms and received either a big drop of juice (~1ml) or a small drop of juice (~0.2ml).

Each day included multiple blocks of trials. Each block contained 28 trials; 1/3 of which were forced information, 1/3 forced random, and 1/3 free choice. Trial types were pseudo-randomly interleaved to counterbalance trial type and target location. After the monkeys had learned to show a stable preference, the identities of the colors were permanently reversed in order to exclude the possibility that the observed preference was driven by the colors of the stimuli.

### Social information-seeking task

In order to develop an abstract version of this advanced information-seeking paradigm, we leveraged monkeys’ natural interest in social information depicted on face. We aimed to develop a social version of this information-seeking paradigm that was as comparable to the juice size information-seeking task as much as possible (**Fig. 2**). Rhesus monkeys are highly sensitive to social images of conspecific faces (Mendelson et al., 1982; Guo et al., 2003; Shepherd et al., 2006; Dal Monte et al., 2016, 2022). In the advanced social information-seeking paradigm, the juice magnitude was always fixed. Instead, the monkeys could choose a cue that tells them in advance the valence, or facial expression, of a monkey face that later appeared in each trial.

The monkeys initiated a trial and made a choice between revealing advanced social information and having random information by fixating on one of the two occluders in the same fashion as the advanced juice size information-seeking task. After the animal fixated on the occluder, they were again given 2,000ms to view the cues. These cues were either informative or random. However, the shape of these cues now predicted the valence of an upcoming picture of a monkey face. After the cue disappeared, a face appeared in the center of the screen for 2,000ms. Monkeys were not required to look at the face. The picture could be either a threatening face or a neutral face. A pool of 88 faces for each valence was used over the course of the experiment, with 8 faces in each category per day. After the face disappeared, the monkeys entered a reward state for 300ms, where they always received a medium-sized pulse of juice (~0.6ml). We used the same block design as in the juice information-seeking task described above.

### Data Analysis

All choice data were recorded and analyzed in MATLAB. Gaze data were extracted from Eyelink data files. Preference for informative versus random cues in both the juice size and social information-seeking tasks was analyzed using a two-tailed t-test for each stimulus pairing.

To measure the looking behavior toward the cues or the faces, both choice and forced trials were analyzed. All looking events to the stimulus of interest counted toward total looking time. Gaze data were normalized to account for individual differences in looking behavior by dividing looking duration during the analysis epoch by the mean looking duration per monkey over all trials for each category of stimuli, cues, or faces. Looking duration was analyzed via ANOVA with the trial outcome (information or random) and trial valence (threat or neutral) as factors. Looking duration to the cues in the juice size information-seeking task was analyzed using a two-way ANOVA with the outcome (information or random) and task type (juice size or social) as factors.

## Results

### Monkeys prefer to seek advanced social information

We first replicated Bromberg-Martin and Hikosaka’s original finding that monkeys prefer to seek advanced information about juice reward magnitude (**Fig. 3**) in the nonsocial, juice size information-seeking task. M1 chose advanced information on more than 95% (±13% s.d. across days) of choices in the original stimulus mapping (t_(770)_=56.5, p<0.001, two-tailed t-test) (**Fig. 3A–B**). After the colors of the informative and random cues were reversed, M1 still preferentially chose to see advanced information on more than 90% (±20% s.d.) of choices (t_(446)_=68, p<0.001). Similarly, M2 chose advanced information on more than 80% (±17% s.d.) of choices in the original stimulus mapping (t_(306)_=38.2, p<0.001) and about 60% (±24% s.d.) of choices after the stimulus reversal (t_(406)_=8.12, p<0.001) (**Fig. 3C–D**). We also observed individual differences between the two monkeys trained on the juice size information-seeking task (on average 80–100% across monkeys), similar to Bromberg-Martin and Hikosaka’s original finding. Therefore, we first replicated advanced information-seeking preferences in our two monkeys.

**Figure 3.**
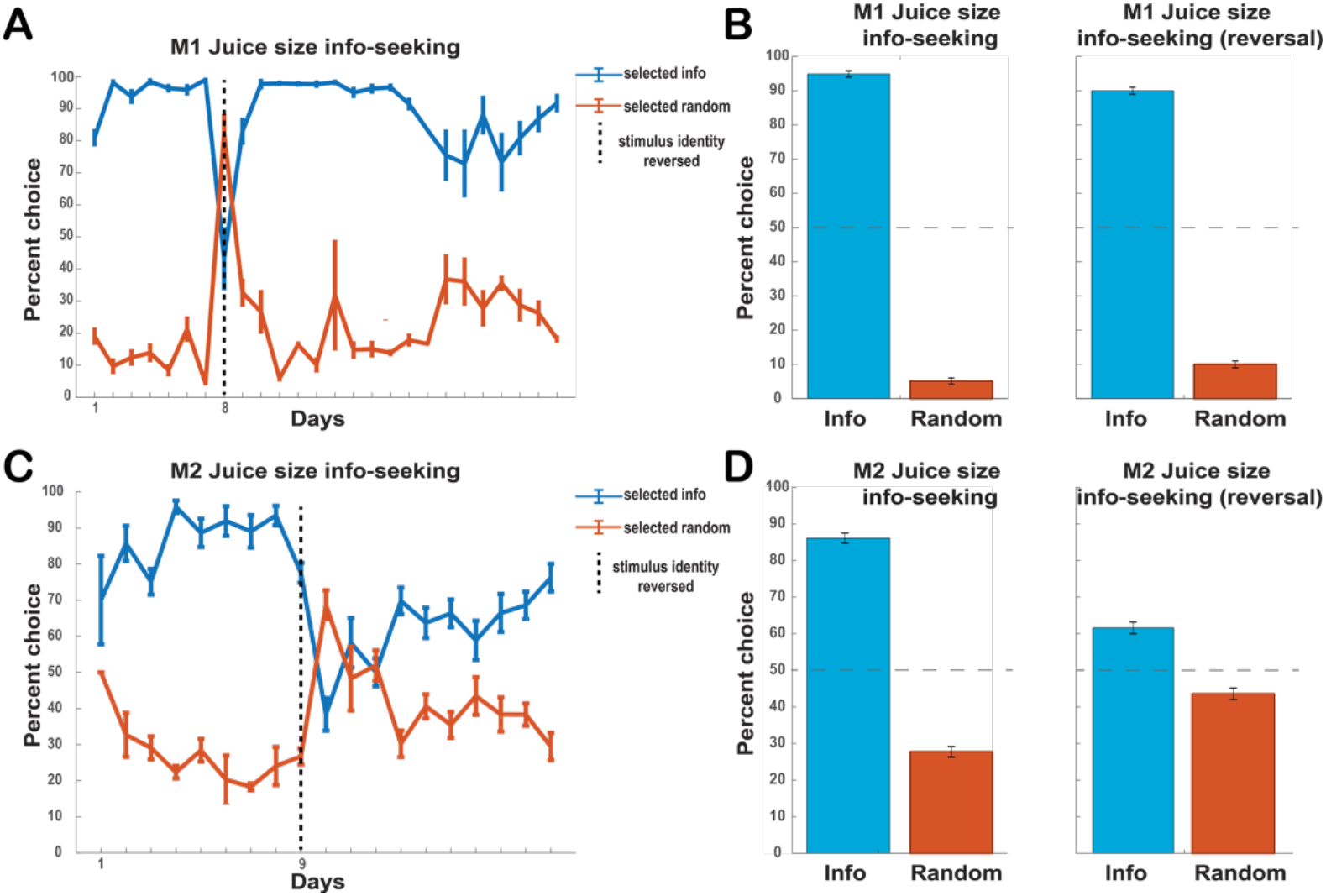
Learning and preference for information-seeking in the juice size information-seeking task. **A**. Percent choice of informative versus random cues across days for monkey 1 (M1), with the date of the reversal of stimuli mapping indicated in the dotted line. **B**. Mean (±s.e.m.) choice percentage across all days for the original stimulus mapping (left) and stimulus reversal (right) for M1. **C–D**. Same format as **A–B** but for monkey 2 (M2). Each monkey demonstrated a preference for advanced information-seeking.

Critically, in the social advanced information-seeking task, both monkeys also showed a preference for advanced information about the valence (threat or neutral facial expression) of the upcoming face (**Fig. 4**). Data shown in **Fig. 4** include learning, as there was no additional training between the juice size advanced information-seeking task and the social advanced information-seeking task. M1 chose advanced social information on 70% (±23% s.d. across days) of choices (t_(474)_=18.7, p<0.001, two-tailed t-test) in the first stimulus mapping (**Fig. 4A–B**). M1 was also trained on a reversed stimulus set (**Fig. 4C–D**) to ensure that M1 was not showing a simple color preference. On the second stimulus set, M1 chose advanced social information on 70% (±18% s.d.) of choices (t_(60)_=8.7, p<0.001). Similarly, M2 chose advanced social information on 60% (±22.8) of choices (t_(136)_=5.5, p<0.01) after learning (**Fig. 4 E–F**). We did not test M2 on a reversed stimulus set as M2 took multiple days to develop this preference, making it less likely that M2 was simply expressing a color preference.

**Figure 4.**
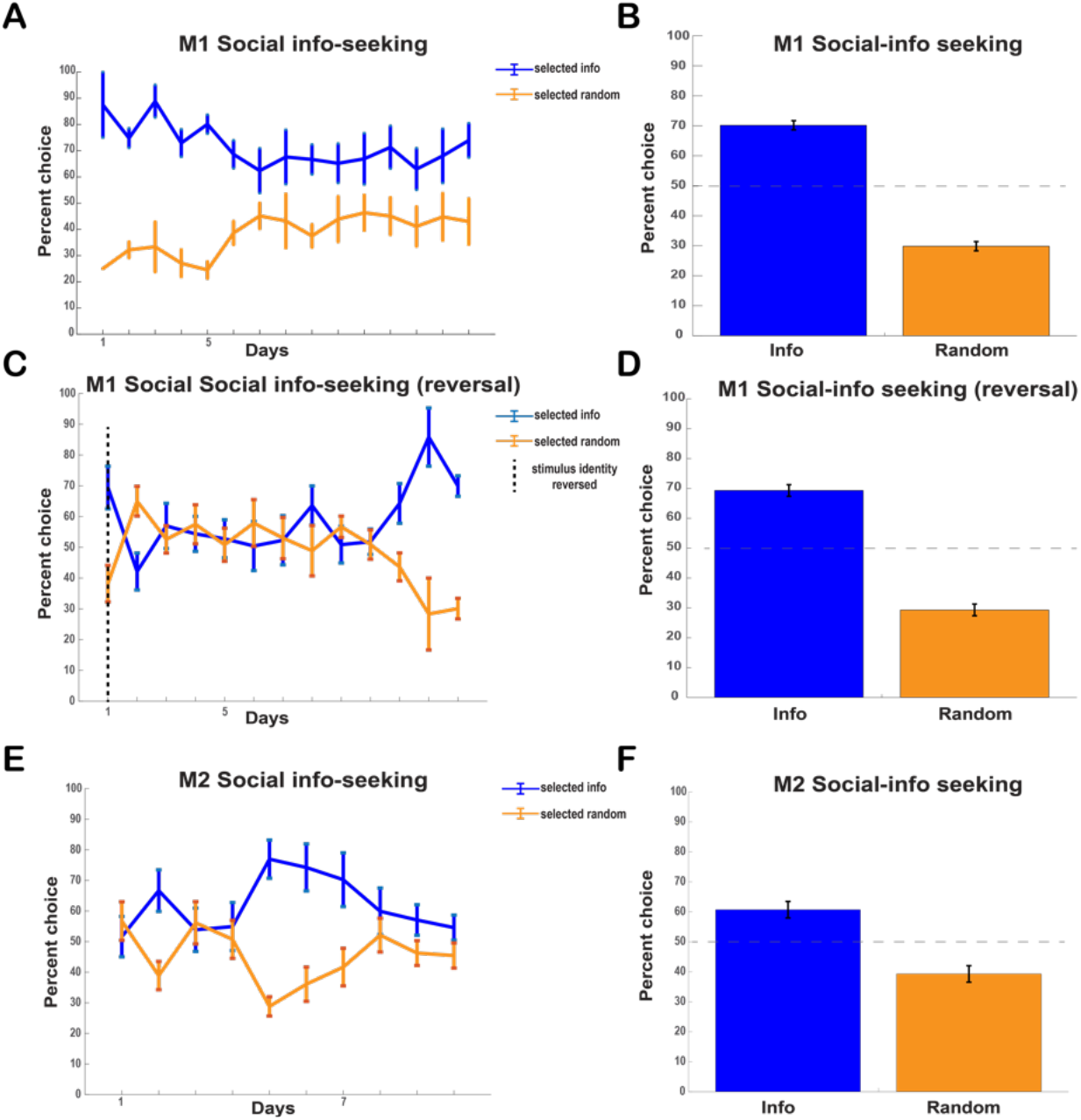
Learning and preference for information-seeking in the social information-seeking task. **A**. Percent choice of informative versus random cues across days for monkey 1 (M1). **B**. Mean (±s.e.m.) choice percentage across all days for the original stimulus mapping for M1. **C**. Percent choice of informative and random cues across days for M1 following the reversal of stimulus mapping (the dotted line as the day of the reversal). **D**. Mean (±s.e.m.) choice percentage across all days for the reversed stimulus mapping for M1. **E–F**. Same format as **A–B** for monkey 2 (M2). Each monkey showed a preference for advanced social information-seeking.

Overall, monkeys preferred advanced information in the advanced social information-seeking task similarly to the juice size information-seeking task. Thus, monkeys were also motivated to seek social information, suggesting that information-seeking preference shown by Bromberg-Martin and Hikosaka can be generalizable to other categories of information. This generalized preference is consistent with the possibility that information, including social information, is inherently rewarding to monkeys.

Interestingly, there was a difference in the overall information-seeking percentage between the juice size information-seeking and the social information-seeking tasks. While monkeys were initially trained on the juice size task and subsequently trained on the social task, later data collection of the two tasks occurred on alternating days, making it unlikely that the lower overall level of preference in the social task simply reflected a decline in general information-seeking behaviors over time. Rather, it is more likely that this difference in the level of information-seeking reflects the additional cognitive load of social stimulus complexity or the diminishing importance of social information as it becomes less immediately self-relevant in this task. Alternatively, it may be driven by the fact that seeking advanced information about primary consumable rewards is simply more motivating compared to seeking information about abstract classes of information, even though more abstract information still has inherent value.

### Gaze behaviors reveal additional insights into seeking advanced social information

While the primary purpose of this task was to assess behavioral preference for advanced social information-seeking, we were also able to examine the monkeys’ gaze behaviors after they made their choices. It is important to note that such looking behavior does not necessarily reflect preference. While looking duration is often used as a proxy for preference (Fujita and Watanabe, 1995; Waitt et al., 2003), it can be an assumption, particularly when that looking behavior is nested within a preference task. However, analyzing this looking behavior may still provide additional insight into how monkeys conceptualize different types of information.

To analyze looking behavior to task-relevant stimuli, we collapsed across choice and forced trials. All of the looking time data shown here used looking duration as the primary measure. The number of fixations was tightly correlated with looking duration so not separately included.

Monkeys looked at the faces about 35% (±16% s.d.) of trials, while they looked at the cues more than half of the times (**Fig. 5A**). As monkeys did not look at the faces very often, it is possible that the pictures of faces were not rewarding in the same way as a reinforcer like juice. While we did not measure the quantity of juice that the monkeys may have failed to consume, monkeys almost always consumed all of the juice rewards available in both the juice size and the social tasks. Additionally, monkeys looked at threat faces for a shorter amount of time than neutral faces, though this effect was only marginally significant (t_(20242)_=-1.7, p=0.07).

**Figure 5.**
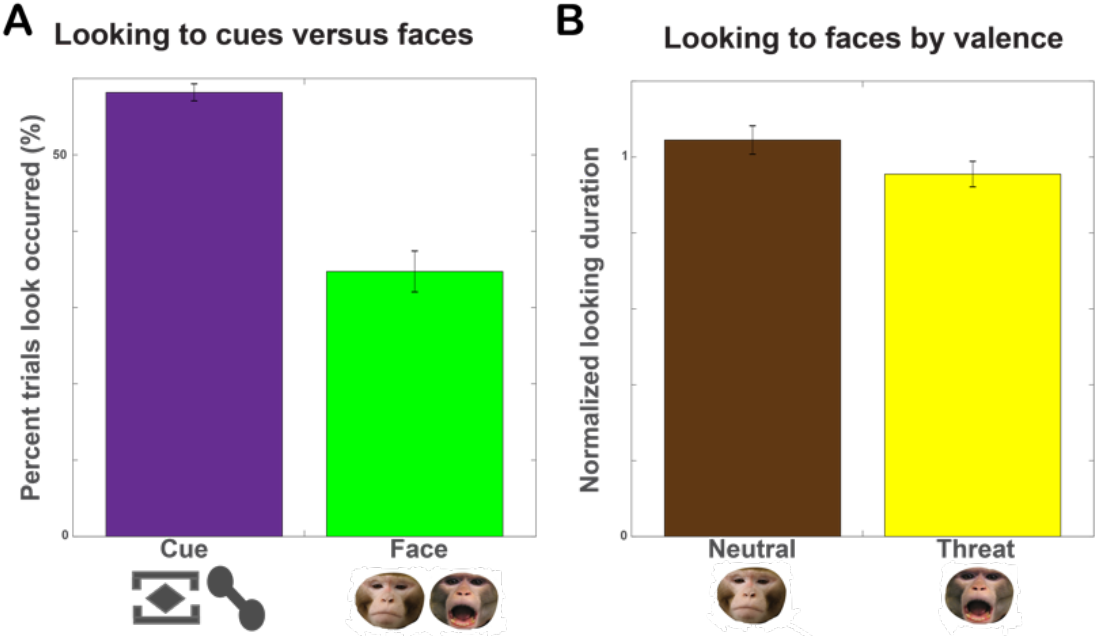
Looking to faces and cues. Monkeys looked at faces on a subset of trials and showed a weak preference in looking duration to one type of facial expression. **A**. Percentage of trials in which monkeys looked to the cues and the percentage of trials in which monkeys looked to the faces. Monkeys did not look at the cues or faces on each trial but were more likely to look at the cues than the faces. **B**. Normalized looking duration to neutral faces and threat faces in the social information-seeking task. There was only a marginal difference in overall looking duration to neutral and threat faces.

Interestingly, splitting the looking duration to the faces by trial outcome (i.e., informative or random) as well as the valence of the picture (i.e., neutral or threat) showed a clearer difference between looking duration to the faces for different outcomes (**Fig. 6A**). A two way ANOVA with outcome and valence (**Table 1**) as factors showed that there was a main effect of the outcome (F_(1,6747)_=11.9, p<0.001). Monkeys looked longer to faces on random trials (1.1±1.6) than on informative trials (0.9±1.5), indicating that monkeys looked more at the face if they did not receive information about what valence it would be in advance. If monkeys wanted to know the valence of the faces but did not receive advanced information, they looked at the faces longer, suggesting that monkeys may have been attempting to use advanced information about the valence of the face to avoid the effort or need to look at the faces. Moreover, there was an interaction between outcome and valence (F_(1,6747)_= 8.73, p=0.05), in which monkeys looked slightly more to threat faces on random trials than threat faces on informative trials as well as slightly less to threat faces on informative trials than to neutral faces on informative trials.

**Figure 6.**
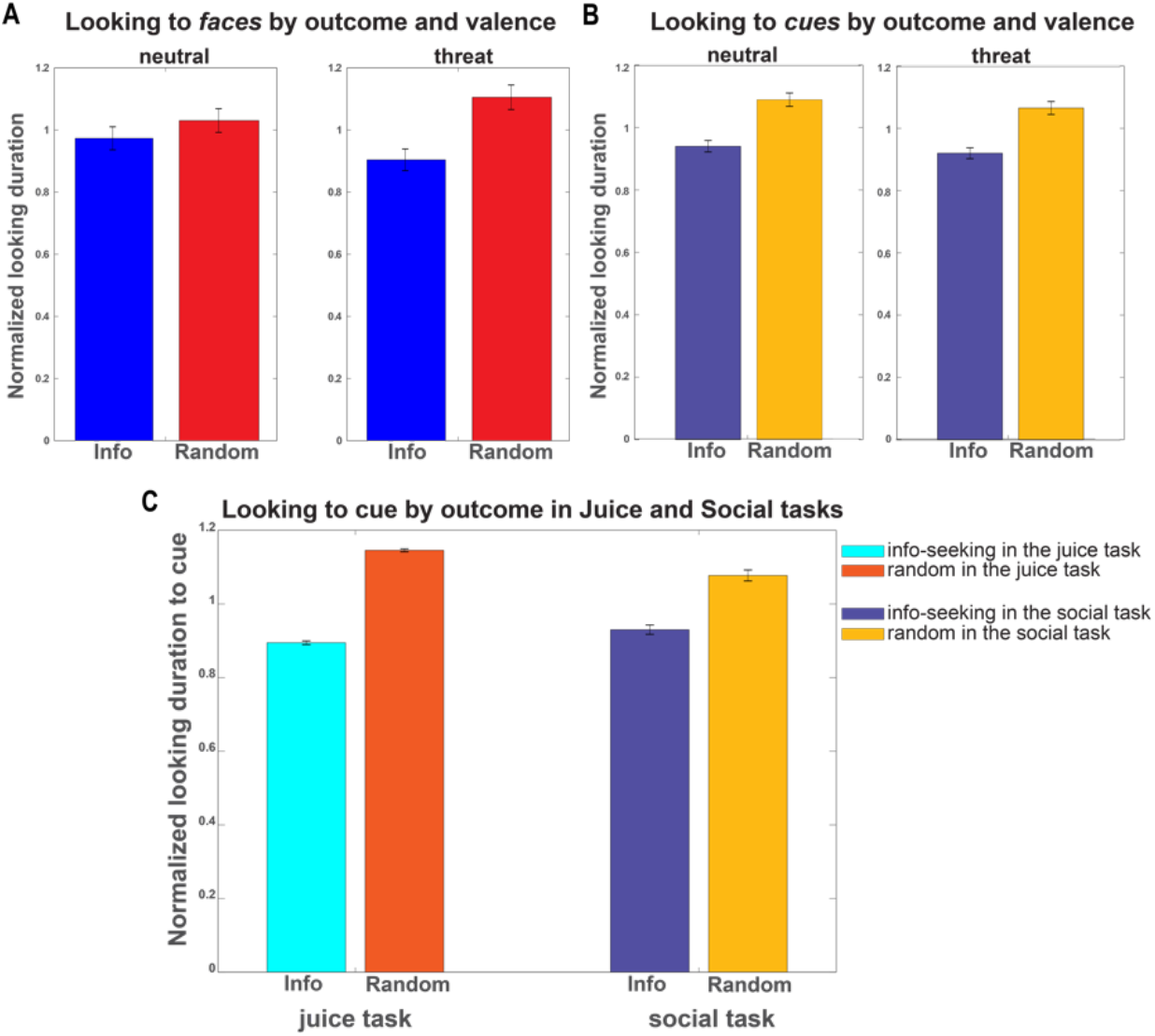
Looking to faces and cues. Monkeys attend to faces and cues more on random trials. **A**. Monkeys looked more to the face on random trials than on informative trials. **B**. Monkeys looked more at the random cues than the informative cues. **C**. Monkeys look more at the random cues than the informative cues in both the juice task and the social task.

**Table 1–3.**
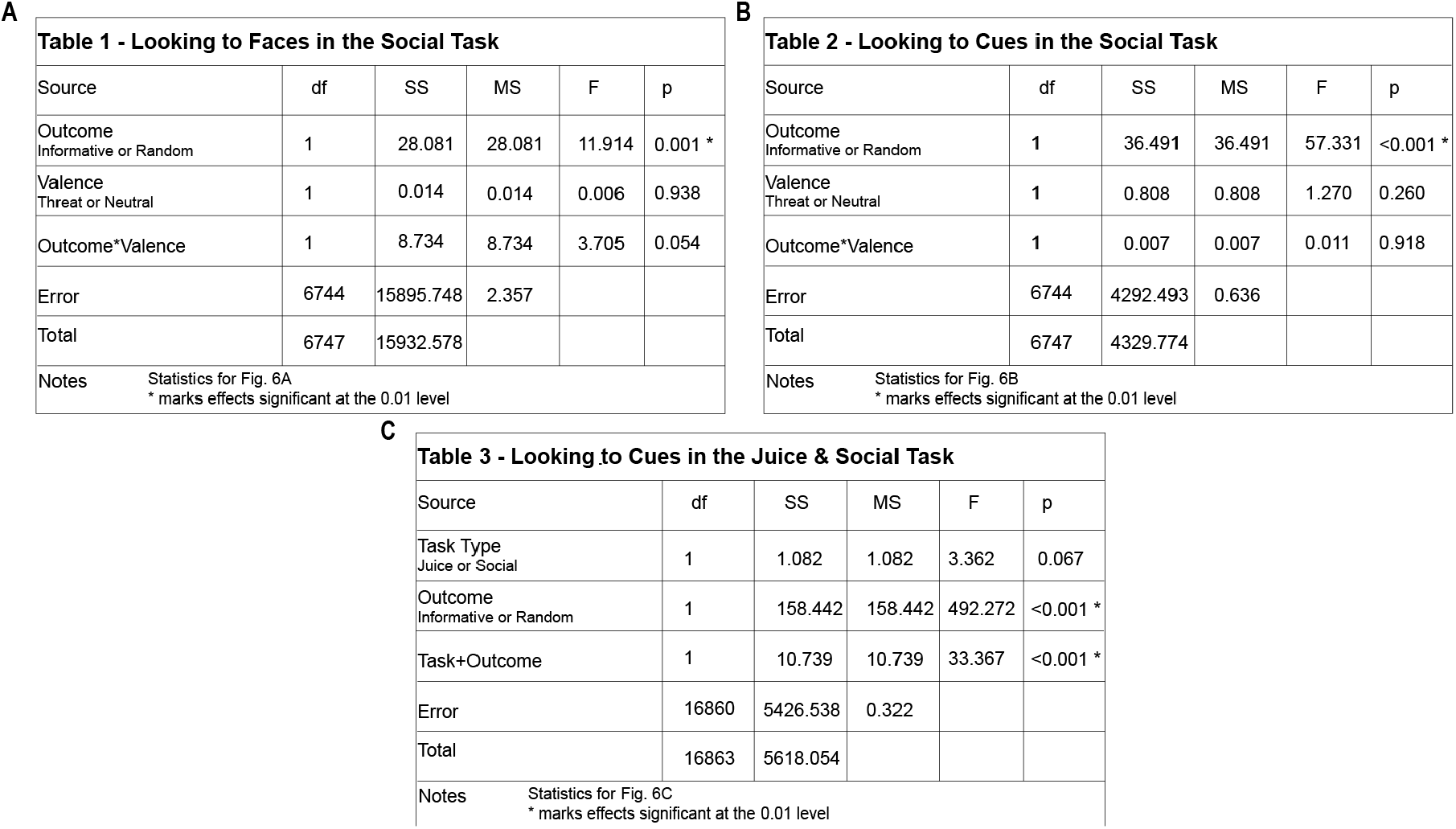
Statistical comparisons for Figure 6. **A**. Table 1. **B**. Table 2. **C**. Table 3. See texts.

|  |  |  |  |  |  |  |  |  |  |  |  |
| --- | --- | --- | --- | --- | --- | --- | --- | --- | --- | --- | --- |
| **A** |  |  |  |  |  | **B** |  |  |  |  |  |
| **Table 1 - Looking to Faces in the Social Task** |  |  |  |  |  | **Table 2 - Looking to Cues in the Social Task** |  |  |  |  |  |
| Source | df | SS | MS | F | p | Source | df | SS | MS | F | p |
| Outcome Informative or Random | 1 | 28.081 | 28.081 | 11.914 | 0.001 * | Outcome Informative or Random | 1 | 36.491 | 36.491 | 57.331 | <0.001 * |
| Valence Threat or Neutral | 1 | 0.014 | 0.014 | 0.006 | 0.938 | Valence Threat or Neutral | 1 | 0.808 | 0.808 | 1.270 | 0.260 |
| Outcome*Valence | 1 | 8.734 | 8.734 | 3.705 | 0.054 | Outcome*Valence | 1 | 0.007 | 0.007 | 0.011 | 0.918 |
| Error | 6744 | 15895.748 | 2.357 |  |  | Error | 6744 | 4292.493 | 0.636 |  |  |
| Total | 6747 | 15932.578 |  |  |  | Total | 6747 | 4329.774 |  |  |  |
| Notes Statistics for Fig. 6A<br>* marks effects significant at the 0.01 level |  |  |  |  |  | Notes Statistics for Fig. 6B<br>* marks effects significant at the 0.01 level |  |  |  |  |  |
| **C** |  |  |  |  |  |  |  |  |  |  |  |
| **Table 3 - Looking to Cues in the Juice & Social Task** |  |  |  |  |  |  |  |  |  |  |  |
| Source | df | SS | MS | F | p |  |  |  |  |  |  |
| Task Type Juice or Social | 1 | 1.082 | 1.082 | 3.362 | 0.067 |  |  |  |  |  |  |
| Outcome Informative or Random | 1 | 158.442 | 158.442 | 492.272 | <0.001 * |  |  |  |  |  |  |
| Task+Outcome | 1 | 10.739 | 10.739 | 33.367 | <0.001 * |  |  |  |  |  |  |
| Error | 16860 | 5426.538 | 0.322 |  |  |  |  |  |  |  |  |
| Total | 16863 | 5618.054 |  |  |  |  |  |  |  |  |  |
| Notes Statistics for Fig. 6C<br>* marks effects significant at the 0.01 level |  |  |  |  |  |  |  |  |  |  |  |

In our tasks, the informative cue indicated information about the upcoming stimulus. Therefore, we expected monkeys to attend to the cue even though it was optional given their choice preference for receiving advanced information. Monkeys looked at the cue on slightly more than half of the trials (**Fig. 5A**). This might reflect that monkeys learned the meaning of the cues rapidly and may be able to peripherally view them without having to fixate directly. Monkeys also attended to the cues more than to the faces (t_(66)_=8.09, P<0.001) (**Fig. 5A**).

We additionally hypothesized that monkeys would look longer towards the informative cue as it could take on the characteristics of a secondary reinforcer by association with the upcoming stimuli. However, surprisingly, monkeys looked longer to the random cue than the informative cue (**Fig. 6B**). A two-way ANOVA with outcome and valence as factors (**Table 2**) found a main effect of outcome (F_(1,6747)_=57.3, p<0.001), where looking duration was longer to the random cues (1.08±0.84, normalized looking duration) than informative cues (0.93±0.75) (p<0.001). This was true regardless of the valence of the upcoming picture; both monkeys looked longer to the random cue on threat and neutral trials. To assess if this is a general tendency in an advanced information-seeking framework, we compared looking duration to the cues predicting the valence of the faces in the social information-seeking task with looking duration to the cues predicting juice size in the juice size information-seeking task. We found that this pattern was preserved (**Fig. 6C**). That is, similar to the results from the social information-seeking task, monkeys looked longer to the random cues in the juice information-seeking task (1.1±0.3) than to the informative cues in the juice size task (0.9±0.4, p<0.001). A two-way ANOVA with task type (juice or social task) and outcome (informative or random) (**Table 3**) as factors found a main effect of outcome (F_(1,16863)_=492.3, p<0.001) and interaction between task type and outcome (F_(1,16863)_=33.4, p<0.001). Over both task contexts, monkeys looked at the random cues more than the informative cue, even though the random cue was not predictive, suggesting that randomness may have an attention-capturing property in the context of seeking advanced information. Furthermore, this finding suggests that the cues do not simply take on the associated value of what they predict in the advanced information-seeking task as secondary reinforcers.

It is important to note that monkeys performed the juice size and social information-seeking tasks on separate days. However, all comparisons between looking duration to informative and random cues in the juice task and informative and random cues in the social task were significant. While monkeys looked longer to the random cues in both tasks, they looked longer to the random cue in the juice size task than they looked to the random cue in the social task. Monkeys also looked longer to the informative cue in the social task than they did to the informative cue in the juice size task. These interesting relationships between looking behaviors in the context of nonsocial and social information-seeking behaviors warrant further investigation.

## Discussion

We first demonstrated that the preference for seeking advanced information in the juice-size information-seeking task is replicable and robust in monkeys. Critically, we extended this finding to show that a preference for advanced information can be effectively found in the realm of social information-seeking as well, indicating that monkeys chose to seek information about abstract categories.

In early studies of curiosity (Berlyne, 1966), it was suggested that animals do not have epistemic curiosity or curiosity that serves to satisfy an intrinsic desire for knowledge. Our results provide an insight into curiosity in nonhuman primates. The preference for choosing advanced information about abstract categories is consistent with the notion that monkeys prefer to know for the sake of knowing. The choice preferences in both types of advanced information-seeking tasks observed here may be part of a generalized curiosity for the information itself, reflecting a desire to know for the sake of knowing. Social curiosity is strongly present in humans. This curiosity may therefore originate from a preference for seeking information found in nonhuman primates and perhaps also in other animals living in groups that have social structures with predictable social relationships amongst the members.

While the monkeys in this task showed a clear preference for advanced information, we examined their gaze behaviors during both task types to further explore whether or not the information is inherently rewarding. A key dilemma in studying gaze patterns is that the interpretation can be uncertain with regard to cognitive operations. Longer looking duration is often postulated to be related to a surprising event (Baillargeon et al., 2010) or preference (Fujita and Watanabe, 1995; Waitt et al., 2003). As our task was primarily designed as a behavioral choice preference task, we cannot isolate exactly what was driving the looking duration effects.

With that caveat in mind, monkeys looked differently, albeit marginally, at threat and neutral faces in the advanced social information-seeking task. More interestingly, monkeys looked longer at the faces on random trials where they did not get advanced information about facial expressions. This suggests that, when presented with a non-predictive cue, they were more inclined to view the faces longer to process their valence and receive the information. It also raises questions about why the monkeys viewed the faces at all, and if their advanced information-seeking preference was providing some utility. It is also possible that monkeys may have used the advanced information to avoid having to look at the faces. If so, this could have been a potential confound in the social advanced information-seeking task – that is, these monkeys may have been able to utilize the advanced information adaptively. In order to isolate “pure” curiosity in the future, passively received stimuli like auditory tones may be more useful as they will be harder to be exploited in an experimental setting.

In our paradigm, the monkeys also looked relatively longer at the random cues than the informative cues. While monkeys preferred advanced information about what social category an upcoming stimulus would belong to, they did not prefer to look longer at the cues that carried that advanced information. The fact that monkeys looked longer at the random cue suggests that the cues did not simply take on the characteristics of a secondary reinforcer. It is possible that the amount of looking at the informative cues was sufficient to extract the relevant information. Furthermore, the random cues may have additional value from an unanticipated source, like exploration or attraction to risk (Heilbronner and Hayden, 2013). That is, longer looking duration at the random cues might be a manifestation of the attraction that uncertainty holds, particularly in a learning environment (Platt and Huettel, 2008). Additionally, randomness could be more salient than predictiveness.

Overall, our results support that monkeys are not only motivated to seek information about primary food rewards, but also seek information about more abstract, social information. As the monkeys in the advanced social information-seeking task chose to seek information about categories that reflected an abstract idea – threat or neutrality – it is possible that monkeys will also demonstrate curiosity about a wide variety of abstractions. If information itself is inherently motivating, this preference should extend to increasingly non-egocentric concepts, such as basic stimulus attributes like color, pitch, or location. If further experiments find that monkeys truly value any type of information, it raises radical questions about the nature of curiosity itself. Rather than representing a high-level impetuous to learn, curiosity may represent a preference for certainty. Whether or not curiosity is differentiable from a preference for predictability will require further experimentation.

### Future directions

Our findings suggest that it is likely that monkeys will show a preference for advanced information about even more abstract categories of stimuli than faces, such as the visual characteristics of shapes. Further investigations of this topic could focus on visual stimuli and exploit categories such as color, geometry, or pattern. Alternatively, other sensory modalities could be studied. While visual stimuli can be attended to or avoided, auditory stimuli must be involuntarily received by the animal, even more so than primary rewards like juice. Testing different modalities of curiosity could further help divorce information-seeking from both reinforcement and utilization.

Exploring the limits of what kinds of information are valuable to organisms, and why, will help determine if curiosity truly functions as an evolved, generalized drive state. It is possible that generalized curiosity is more common in animals that live in complex social environments, as secondary reinforcers become critical information for survival when they are generated by the members of social groups. Additionally, it is possible that curiosity as a drive state is derived from foraging for primary reward, and that it is shared by all organisms that strive to exploit their environment for survival. If monkeys show a preference for advanced information about stimuli that are practically irrelevant to themselves, it will even more strongly indicate that they have a general drive for information. Humans describe the desire to know just for the sake of knowing. This drive for curiosity may be mediated by an internal drive that is not unique to humans.

## Author Contributions

S.W.C.C. and J.A.J. designed the study, and S.W.C.C. and J.J. wrote the paper. J.J. performed the experiments. J.J., N.A.F., and S.W.C.C. analyzed the data.

## Acknowledgments

This work was supported by Alfred P. Sloan Foundation (FG-2015-66028).

